# Chloroplast-mitochondria synergy modulates responses to iron limitation in two *Thalassiosira* diatom species

**DOI:** 10.1101/2025.11.28.691171

**Authors:** Jhoanell Angulo, Clarisse Uwizeye, Pascal Albanese, Mathilde Menneteau, Stéphane Ravanel, Pierre-Henri Jouneau, Giovanni Finazzi, Florence Courtois

## Abstract

Iron is naturally present at low levels in modern oceans, but it remains essential for marine life. Ocean-dwelling organisms such as oceanic phytoplankton must therefore adapt to the available levels. Among phytoplankton, diatoms are a highly diverse and successful taxon that includes the *Thalassiosira* genus. As a group, diatoms contribute around 20% of global primary photosynthetic productivity, they have also developed specific resources allowing them to thrive in low-iron regions. However, the major biological factors underlying their success in these ocean environments remain unknown. Here, we compared two *Thalassiosira* species: *T. oceanica* from iron-poor open-ocean; and *T. pseudonana*, from iron-rich coastal waters. Since iron is essential for both photosynthesis and respiration, we examined the specificities of the bioenergetic machineries in these organisms using a combination of photo-physiological, proteomics, and FIB-SEM methods. We particularly focused on chloroplast-mitochondrial coupling, a mechanism deployed by diatoms to ensure optimal transfer of photosynthetic products to promote cell growth. This study of this mechanism in the context of iron limitation reveals that the two diatoms differentially remodel chloroplast compartments in response to iron limitation. Their tolerance to these conditions is also linked to distinct constitutive mitochondrial architectures.

## Introduction

Diatoms, unicellular photosynthetic marine heterokonts, are major contributors to the oceanic food chain and to carbon dioxide storage, accounting for about 20% of primary production worldwide [1]. Their genomic and metabolic complexity [2, 1, 3], linked to their origins in secondary endosymbiosis, is characterized by frequent horizontal gene transfers [4, 5]. This complexity provides them with the resilience and adaptability needed to survive in diverse and fluctuating environments, where light intensity may vary [6, 7], darkness may be prolonged [8], mineral nutrient availability may fluctuate [9], and nutrient-stimulated blooms may occur [10]. The growth of diatom species and their distribution between turbid coastal and open-ocean waters often follow dissolved iron gradients [11, 12]. Indeed, low iron bioavailability chronically limits photosynthetic carbon fixation in open-ocean areas [13, 14]. Conversely, iron levels in coastal waters tend to be higher due to supply from the land [15, 16]. To rationalize the ecological success of diatoms in oceans, we need to understand how they adapt and/or acclimatize to distinct environments [17].

Iron-containing proteins – such as iron-sulfur centers, cytochromes – are highly represented in chloroplasts and mitochondria, the organelles responsible for photosynthesis and respiration, respectively [18, 19]. These two processes are thus potentially impacted by iron limitation. The iron-containing proteins act as electron carriers transducing reducing power across cell compartments and contributing to the generation of the transmembrane proton gradient required for bioenergetic processes in the cell. The photosynthetic capacity of phytoplankton tends to be limited by iron restriction [20, 21]. Diatoms have developed mechanisms to allow energetic cross-talk between chloroplasts and mitochondria [22, 23] as a strategy to ensure optimal carbon fixation and cell growth [24, 25, 23]. However, the interactions between the subcelluar compartments and their consequences on both cellular organization and bioenergetic processes have yet to be characterized in the context of iron limitation. Previous proteomics and biochemical data suggest increased coupling between photosynthetic and respiratory functions in diatoms grown under low iron levels [24, 26]. However, this cross-talk has not been fully elucidated, and we lack information on essential physiological and morphological adaptations in the context of iron limitation.

*Thalassiosira oceanica* is a low-iron-tolerant diatom. Its responses to limited iron availability involve specific adaptations, such as constitutive substitution in chloroplasts of iron-containing cytochrome c_6_ by copper-containing plastocyanin [27] and of ferredoxin by flavodoxin [28]. No equivalent changes to mitochondrial proteins have been observed in this species, and the relative abundance of iron-rich mitochondrial complexes is preserved under iron starvation [3]. Other diatoms react similarly to iron starvation, with transcription of iron reductase reported in *Thalassiosira pseudonana* and of flavodoxin in *Thalassiosira weissflogii* [29]. In *Phaedactylum tricornutum,* high-affinity carrier proteins, such as ISIP2a, are expressed to enhance iron supply [30]. In addition to these iron-uptake and -storage adaptations, iron starvation triggers metabolic and physiological responses, such as modified lipid metabolism, increased degradation of cell components [3], as well as recycling and reallocation of carbon and nitrogen [3, 24, 31].

To complete this picture and assess the overall relevance of these responses, we conducted a comparative analysis of laboratory-cultured *T. oceanica* and of its close relative species *T. pseudonana*. The two diatoms were grown in media containing high and low iron concentrations, typical of their respective natural environments [12, 32]. The iron-rich and iron-poor media represented challenging conditions in terms of iron availability for each species. We then combined morphological, physiological, and proteomics approaches to shed light on the strategies deployed by the two organisms to cope with low iron availability. Our data revealed species-specific adaptations of subcellular morphology and physiological acclimation to sustain chloroplast-mitochondrial coupling.

## Results

### Mitochondrial architecture differs between *T. oceanica* and *T. pseudonana*

We first compared cell morphologies in *T. oceanica* and *T. pseudonana* cells grown in an iron-rich medium (Fig. 1, Fig. S1). Using Focused Ion Beam Scanning Electron Microscopy (FIB-SEM), we imaged thin lamellae from fixed cells and derived three-dimensional (3D) structures. This analysis revealed that *T. oceanica* cells are larger than *T. pseudonana* cells (Fig. 1A-C). In both species, the frustule and ribs of the valves were visible at the working voxel size of 16 nm (Fig. 1A, B). All cells contained one large vacuole, occupying approximately 40% of the total cell volume, consistent with [12, 33] (Fig. 1D-F). The proportion of the cell volume occupied by all organelles – vacuole plus chloroplasts, mitochondria, nucleus – was equivalent in the two species, at ∼60% of the total cell volume (Fig. 1F). Chloroplasts and mitochondria were located at the cell periphery (Fig. 1G, H). Although the sub-organelle organization of chloroplasts was equivalent in both diatoms, their mitochondrial architecture was distinct. Thus, the relative volume of the mitochondrial compartment and the mitochondrial surface-to-volume ratio were higher in *T. pseudonana* than in *T. oceanica* (Fig. 1I, L). Moreover, the two species had distinct mitochondrial shapes: spherical in *T. oceanica* (Fig. 1J), and elongated in *T. pseudonana* (Fig. 1K). Nevertheless, the inner mitochondrial structures appeared to be the same in the two diatoms (Fig. 1J, K).

**Figure 1.**
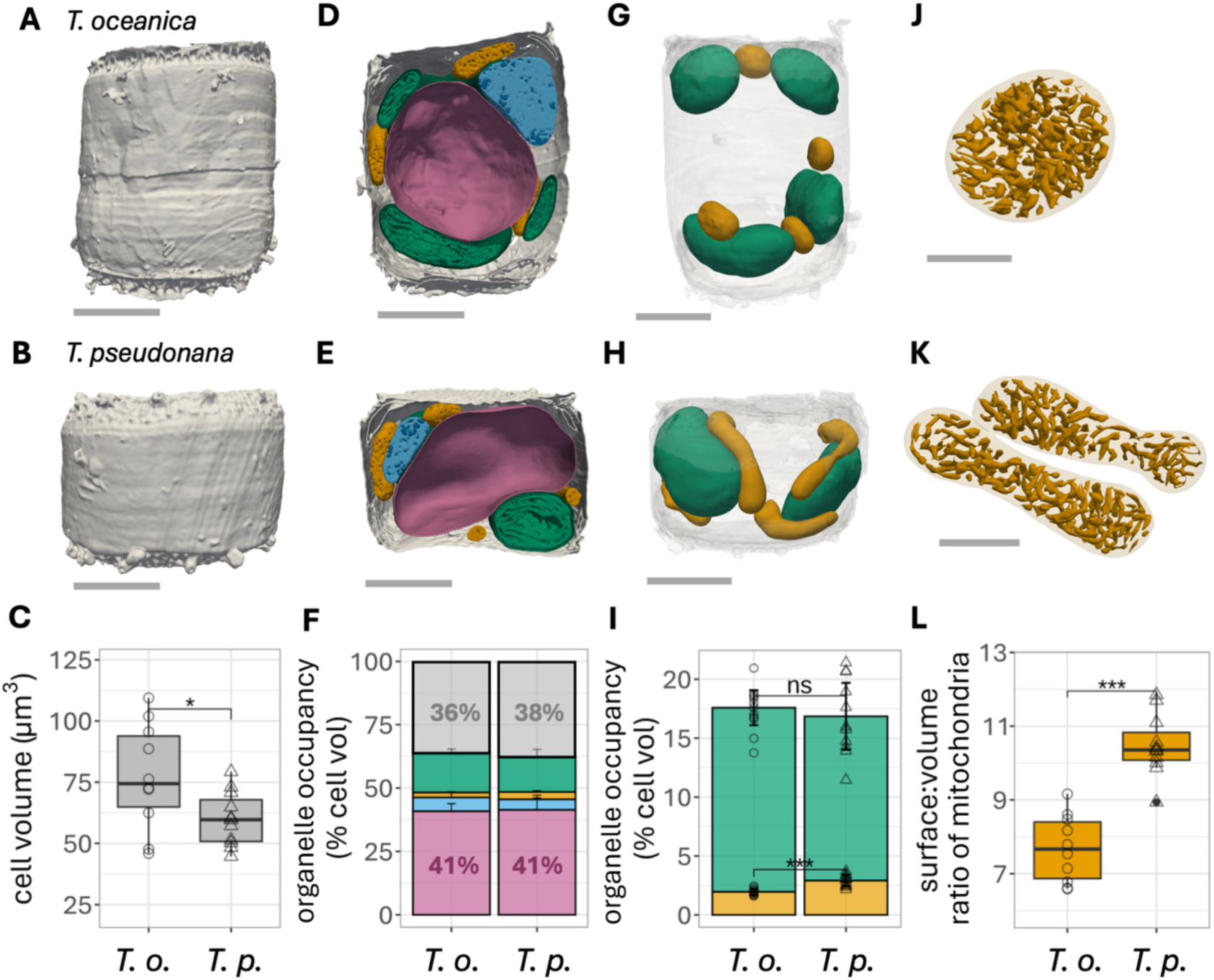
Mitochondrial morphology differs between *T. oceanica* and *T. pseudonana*. *T. oceanica* and *T. pseudonana* cells were observed after growth in an iron-rich medium. The 3D models shown are representative of at least 9 cells from 2-3 independent biological replicates for each condition **(A, B)** 3D scan view of cell morphology. Voxel size = 16 nm. **(C)** Absolute cell volume, derived from quantitative analysis of the 3D models. **(D, E)** 3D models of the main subcellular compartments. Green: chloroplast; yellow: mitochondria; blue: nucleus; pink: vacuole; gray: other cell structures and silica frustule. **(F)** Volumes of the main subcellular compartments, derived from quantitative analysis of occupancy of the 3D models. Error bars: SD. Color code as in D, E. **(G, H)** 3D models highlighting chloroplasts (green), mitochondria (yellow), and cell frustule (gray). **(I)** Total cell volume occupied by chloroplasts (green) and mitochondria (yellow), derived from quantitative analysis of the 3D models. Error bars: SD **(J, K)** Segmentation of mitochondrial and cristae volumes. **(L)** Surface-to-volume ratio of total mitochondrial compartment in a single cell. **(A, B, D, E, G, H)** Scale bar: 2 µm; **(J, K)** Scale bar = 0.5 μm. **(C, F, I, L)** *T.o.: T. oceanica*; open circles; n=10 cells. *T.p*.: *T. pseudonana;* open triangles; n= 11 cells. Statistical analysis based on paired Welch’s t-test; ***: p ≤ 0.001; ns: non-significant.

In addition to differences in shape, the subcellular location of mitochondria differed between *T. pseudonana* – where they formed a network appressed to the chloroplast envelope – and *T. oceanica –* where they were dispersed in the cytoplasm (Fig. 2A, B). Based on this organization, the total chloroplast surface in proximity to mitochondria, calculated at a maximal distance of 30 nm, as in [34], was higher in *T. pseudonana* than in *T. oceanica* (Fig. 2C-F). This distinct subcellular arrangement of the two organelles may affect the physical interactions between chloroplasts and mitochondria.

**Figure 2.**
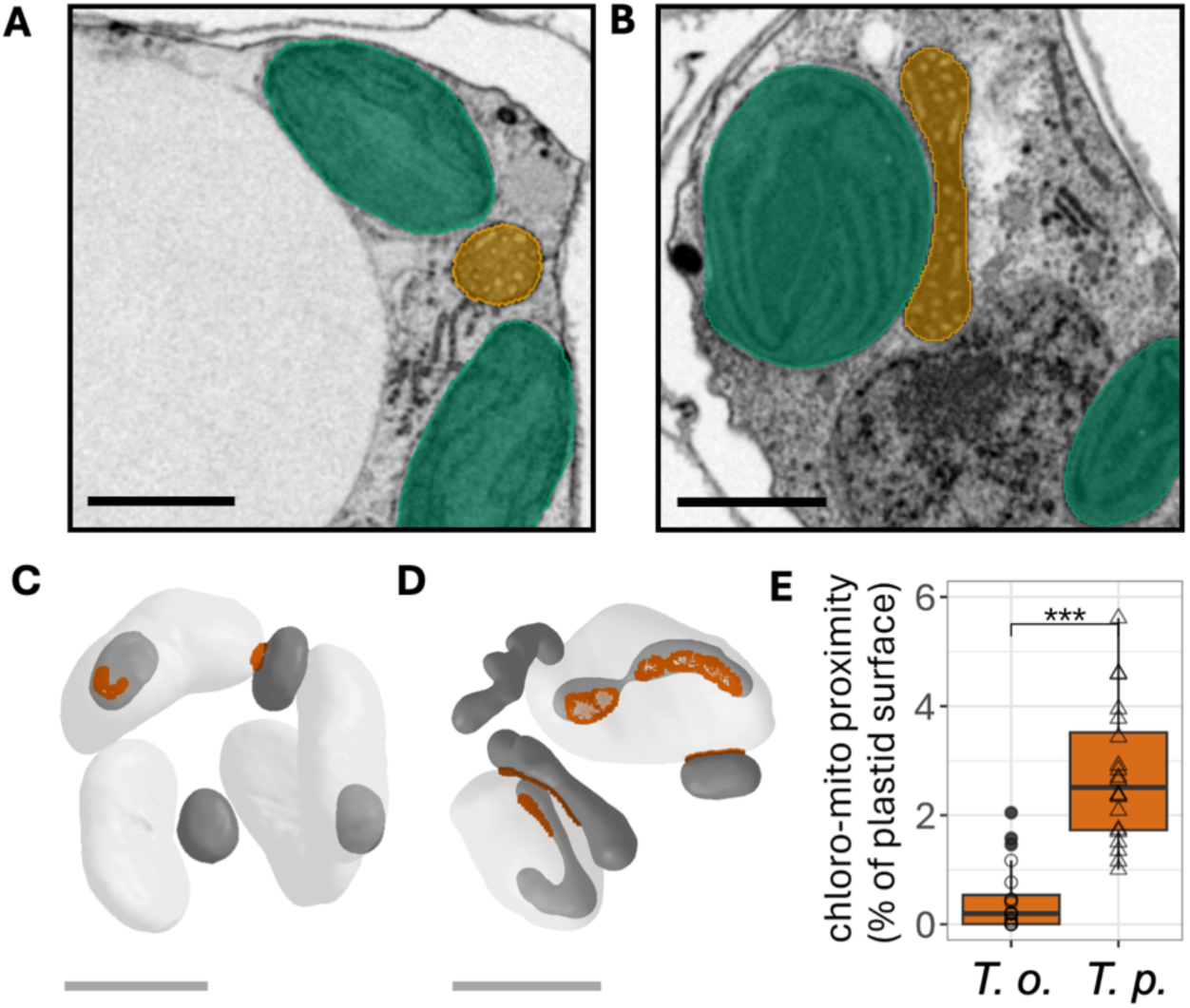
Chloroplast-to-mitochondria proximity differs between *T. oceanica* and *T. pseudonana*. **(A, B)** Sections through cellular 3D volumes, segmented from FIB-SEM micrographs of whole cells, chloroplasts (green), mitochondria (yellow). Scale bar: 1 µm. **(C, D)** 3D models of the cells from A, B showing chloroplasts (light gray) and mitochondria (dark gray), and their proximity (points at ≤ 30 nm are indicated by dark orange shading). Scale bar = 2 µm. Sections and models are representative of at least 9 cells from 2-3 independent biological replicates for each condition. **(F)** Total chloroplast surface at less than 30 nm from mitochondria (%). *T.o.: T. oceanica*; open circles; n= 10 cells. *T.p*.: *T. pseudonana;* open triangles; n= 11 cells. Statistical analysis by paired Welch’s t-test; ***: p ≤ 0.001.

### Iron limitation has distinct effects on cell physiology in *T. oceanica* and *T. pseudonana*

We next compared how the two diatom species cope with differences in iron availability during growth (Table 1). The iron content of cells grown in iron-rich and iron-poor conditions was measured by Inductively Coupled Plasma Mass Spectrometry (ICP-MS), with normalization relative to Mg content (Table 1). A similar trend was observed upon normalization relative to P content (not shown, [35, 36]).

**Table 1.**
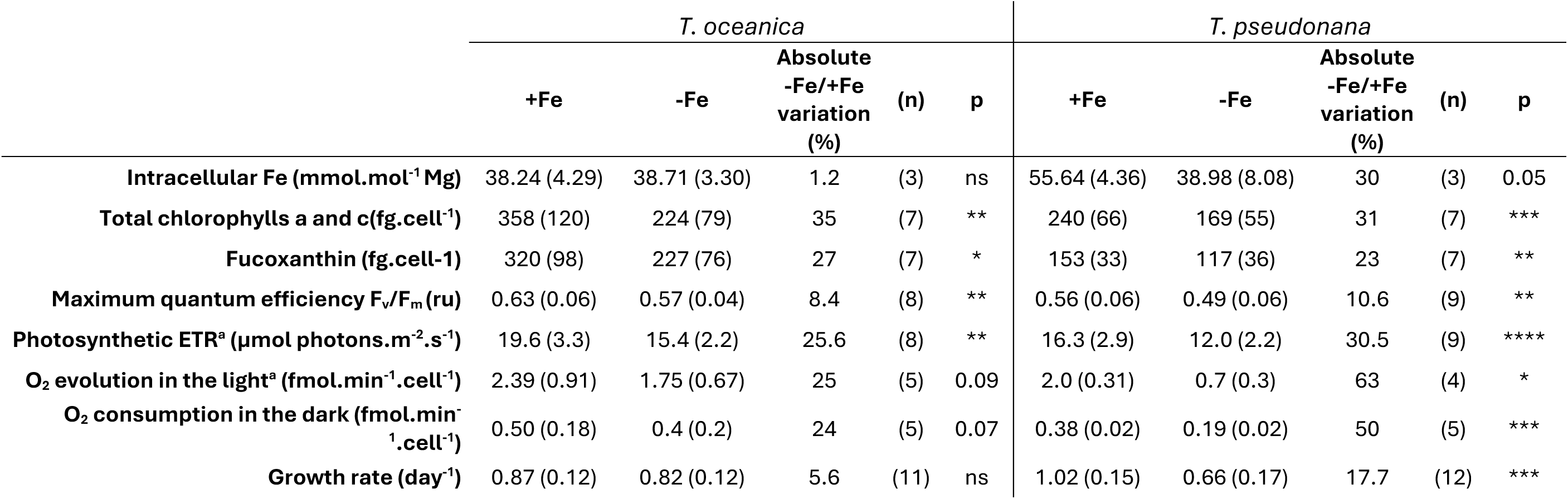
Iron limitation effects on *T. oceanica* and *T. pseudonana* physiology. *T. oceanica* and *T. pseudonana* cells were grown under iron-rich (+Fe) and iron-poor (-Fe) conditions. Absolute -Fe/+Fe variations were calculated as follows: 1 - (-Fe)/(+Fe) * 100. Values in parentheses correspond to SD. (**a**) Light intensity during measurements: 133 µmol photons.m^-2.^s^-1^. (**b**) Light intensity during measurements: 120 µmol photons.m^-2.^s^-1^. (n): number of independent biological replicates (**c**) p: statistical significance as determined by paired Welch’s t-test; ****: p ≤ 0.0001; ***: p ≤ 0.001; **: p ≤ 0.01; *:p ≤ 0.05; ns: non-significant.

In iron-rich conditions, total iron content was significantly higher in *T. pseudonana* compared to *T. oceanica*. In iron-poor conditions, conversely, the total iron content was decreased in *T. pseudonana* cells but not in *T. oceanica* cells (Table 1). These results suggest that the iron uptake capacity of *T. oceanica* does not depend on the iron concentration in the growth medium, at least over the mild range tested in this study, conversely to [32, 37].

In iron-poor conditions, both strains had decreased total chlorophyll and fucoxanthin content compared to the iron-rich condition (∼30% lower in *T. pseudonana* and ∼25% lower in *T. oceanica*; Table 1). The maximum photochemical quantum yield of PSII (Fv/Fm) and the photosynthetic electron transport rate were reduced to a similar extent. These observations reveal the impact of iron limitation on the photosynthetic machinery to be similar in both species. Nevertheless, O_2_ evolution and consumption rates were differently impaired in the two species studied. Thus, in the presence of limited iron, we calculated a 25% reduction in both evolution and consumption in *T. oceanica*, contrasting with a 63% reduction in evolution and a 50% reduction in consumption in *T. pseudonana* (Table 1). Differences between photosynthetic electron flow activity, as determined from the electron transfer rate (ETR) and O₂ evolution measurements, were also observed between species, suggesting possible alternative electron sink.

Finally, in iron-limited conditions, the growth rate of *T. pseudonana* was significantly reduced compared to that of *T. oceanica* (Table 1). Thus, overall, moderate iron limitation led to distinct physiological responses between open-ocean and coastal *Thalassiosira* species.

### In iron-limited *T. oceanica*, functional and protein changes are associated with rearranged organelle morphology

From segmented FIB-SEM micrographs of iron-limited cells, we derived 3D models of chloroplasts and mitochondria, and determined their volumes (Fig. 3). These data revealed differences in chloroplast plasticity between the two species in response to limited iron availability. In *T. oceanica*, the average number of chloroplasts per cell was reduced from 4 to 2 under iron-poor conditions (Fig. 3A, B). The chloroplast-occupied cell volume was also decreased, whereas the volume of the mitochondrial compartment remained unaltered (Fig. 3C). Thus, based on organelle morphology, iron limitation has a more drastic effect on chloroplasts than on mitochondria in *T. oceanica*. In *T. pseudonana,* the number of chloroplasts remained constant at 2 per cell (Fig. 3D, E) and the volume occupancy of chloroplasts and mitochondria (Fig. 3F) were unaffected under iron-poor conditions.

**Figure 3.**
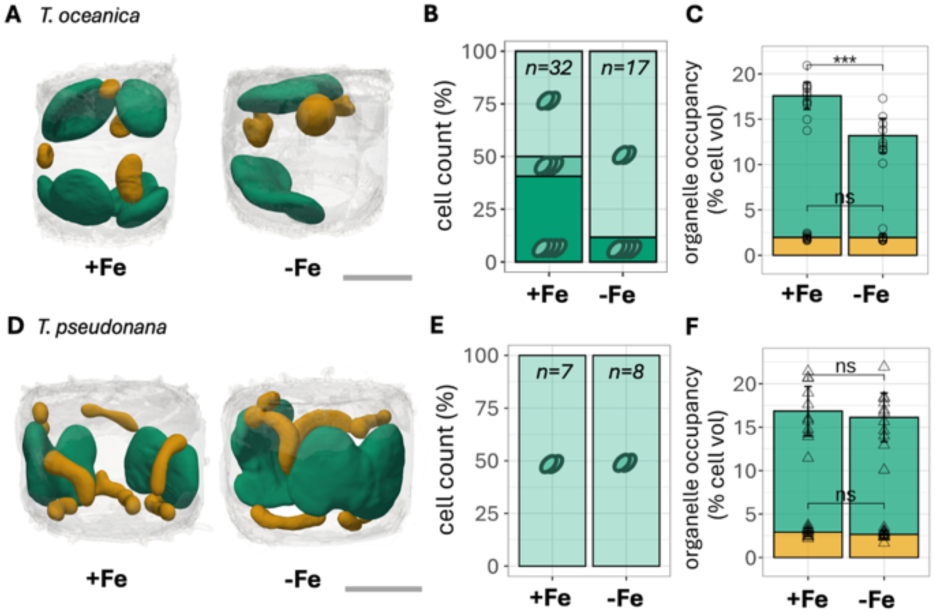
Comparative effects of iron limitation on the subcellular morphology of *T. oceanica* and *T. pseudonana*. **(A, B)** 3D models from segmented FIB-SEM micrographs of chloroplasts (green), mitochondria (yellow). Scale bar 2 µm. 3D models are representative of at least 9 cells from 2-3 independent biological replicates for each condition. **(C, D)** Distribution of cell population by number of chloroplasts per cell under iron-rich (+Fe) and iron-limited (-Fe) conditions. 2 chloroplasts: light green, 3 chloroplasts: medium green, 4 chloroplasts: dark green; n: total number of cells investigated. **(E,F)** Total cell volume occupancy by chloroplasts (green) and mitochondria (yellow), as derived from quantitative analysis of the 3D models. Bars indicate SD. *T.o.: T. oceanica*; circles; n= 10 cells. *T.p*.: *T. pseudonana;* triangles; n= 11 cells. Statistical significance tested using paired Welch’s t-test; ***: p ≤ 0.001; ns: non-significant.

Moreover, the peculiar elongated mitochondrial architecture, and the proximity of chloroplasts and mitochondria were similar in all culture conditions, whatever the iron levels (Fig. S2). The subcellular architecture of the mitochondrial network therefore differs between *T. oceanica* and *T. pseudonana,* and this species-specific feature is independent of iron availability.

To investigate possible whole-cell effects induced by iron limitation in the two cell types, we conducted a comparative proteomics analysis to determine the relative abundance of proteins isolated from cells following growth in iron-poor or iron-rich conditions (Dataset S1, Fig. 4, Fig.S3). Overall, relative abundances of the proteins detected (1628 in *T. oceanica* and 2263 in *T. pseudonana*) were not extensively modified by iron limitation (88% in *T. oceanica* and 96% in *T. pseudonana*). In *T. oceanica*, 73 proteins increased in abundance (4.5% of total identified proteins) and 121 decreased (7.4%) (Fig. S3). In *T. pseudonana*, 30 proteins increased in abundance (1.3%) and 58 decreased (2.6%). To identify function-related patterns in these differentially expressed proteins, we clustered them based on their functions in chloroplasts and mitochondria as described in [38] (Fig. 4). In *T. oceanica,* in line with the physiological changes reported above, most of the iron-limitation-sensitive proteins were chloroplast proteins, with a specific reduction of the Light-Harvesting Complexes (LHC) antenna and of proteins from the photosynthetic Electron Transport Chain (ETC) including subunits of the iron-requiring photosystems I and II, and of cytochromes (Fig. 4). The iron-free plastocyanin electron carrier [39] was detected in *T. oceanica*, but its abundance was unaffected by iron availability during growth (Fig. S3, Dataset S1). Expression of mitochondrial proteins (i.e. iron-rich ETC complexes and Krebs cycle enzymes) tended to be more conserved under iron limitation (Fig. 4). Conversely, in *T. pseudonana*, the relative abundances of most of the photosynthetic ETC and LHC subunits and of mitochondrial ETC complexes were not significantly affected by iron limitation (Fig. 4).

**Figure 4:**
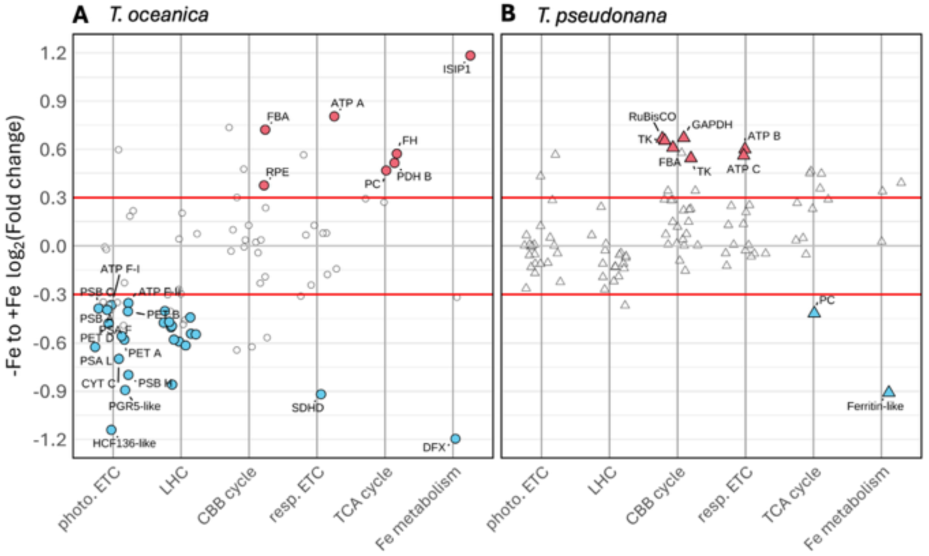
Proteome rearrangement in *T. oceanica* and *T. pseudonana* under iron limitation. The total proteomes of **(A)** *T. oceanica* (circles) and **(B)** *T. pseudonana* (triangles) in iron-poor and iron-rich conditions were analyzed by MS-based label-free quantitative proteomics (Fig. S3). The plot displays the differential abundance (log_2_FC iron-poor vs. iron-rich, y axis) of proteins identified, clustered by their functional categories, as in [38].. Photo ETC: photosynthesis-specific electron transport chain proteins; LHC: Light-harvesting complex proteins; CBB cycle: Calvin-Benson-Bassham cycle proteins; Resp ETC: respiration-specific electron transport chain proteins; TCA cycle: mitochondrial tricarboxylic acid cycle proteins; Fe metabolism: iron-metabolism-related proteins. Blue and red dots highlight proteins that were significantly depleted or enriched, respectively, in samples grown in iron-poor conditions as compared to samples from iron-rich cultures. Thresholds applied were: a log_2_FC cut-off of 0.3 (delimited by horizontal lines on plot) and a P-value cut-off of 0.01 (Limma P-value) for the analysis of proteins clustered by process. Results presented are representative of four independent biological replicates. See Dataset S1 for accession numbers of the proteins listed.

### Photosynthesis and respiration are coupled differently in *T. oceanica* and *T. pseudonana*

The readjustment of the architecture of the chloroplast and mitochondrial networks, and the differences in physiological responses under iron limitation suggest differences in coupling between the photosynthetic and respiratory functions between the two species. We reasoned that, in the presence of mitochondrial dysfunction, the coupling between the organelles might result in reduced photosynthesis, e.g. lower Electron Transfert Rate (ETR). We tested this possibility by exposing cells to a cocktail of mitochondrial inhibitors, including inhibitors of mitochondrial alternative oxidase (salycylhydroxamic acid) and complex III (antimycin A), to target both the cyanide-sensitive and cyanide-insensitive mitochondrial electron transfer pathways [23, 40]. We then measured the photosynthetic ETR, based on chloroplast fluorescence. In iron-rich conditions, in both species, the photosynthetic ETR was reduced in the presence of the respiratory inhibitors (Fig. 5). This suggests that both organisms have mechanisms to ensure metabolic exchange between chloroplasts and mitochondria. However, the effect was more significant in *T. pseudonana* than in *T. oceanica.* When iron availability was limited, mitochondrial inhibitors had an even more drastic effect in *T. pseudonana* (Fig. 5).

**Figure 5.**
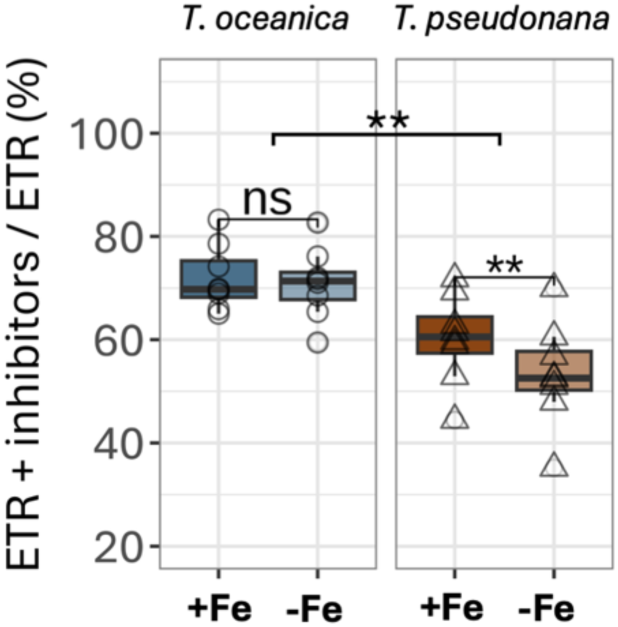
Differences in functional plastid-mitochondria coupling between species could determine tolerance to Fe limitation. Photosynthetic Electron Transport Rate (ETR) estimated from the PSII operating efficiency in the presence of respiratory inhibitors (see Materials and Methods). Light intensity during measurements: 45 μmol.m^-2^.s^-1^ PPFD. Data shown are representative of eight independent biological replicates in each condition; Statistical significance was determined by applying a paired Welch’s t-test; **: p ≤ 0.01; ns: non-significant.

## Discussion

In this study, we compared two closely related diatom species (*T. oceanica* and *T. pseudonana*) that occupy ecological niches distinguished by their iron bioavailability. In laboratory conditions, the two species were grown in media containing iron concentrations reflecting the levels measured at the two sites where the diatoms were isolated. To understand how they acclimatize to the different environments, we characterized their morphology, their physiological responses, and the relative abundances of their proteins under conditions where iron was the only variable parameter.

Morphological assessment revealed distinct organelle architectures in the two diatoms: spherical mitochondria located independently of chloroplasts in *T. oceanica*, contrasting with elongated mitochondria appressed against chloroplasts in *T. pseudonana* (Fig. 1, Fig. 2). These features were unaffected by iron levels in culture media (Fig. 1, Fig. S2), and thus probably represent evolutionary adaptations to their distinct natural habitats (i.e. open-ocean and coastal waters, respectively).

In both diatoms under iron limitation, we observed a very similar sustained chlorosis and decreases in photosynthetic and respiratory rates, although the effect on photosynthetic and respiratory rates was more pronounced in *T. pseudonana* (Table 1). The apparent contradiction between fluorescence- and oxygen-based measurements of photosynthetic activity under iron limitation can be explained by the activity of the plastoquinol terminal oxidase (PTOX), which catalyzes the flow of electrons from plastoquinol to O₂ during the chlororespiration process [41, 42]. This process contributes to the ETR but not to O_2_ evolution, since PTOX converts oxygen to H_2_O in stoichiometric amounts [43]. The increased PTOX activity observed elsewhere [43], was not detected in the proteomics analysis. This apparent discrepancy suggests that, either its accumulation level is below the detection threshold in both species, or the difference in activity is not linked to an increase in protein abundance.

Despite the physiological adaptations observed, the cellular iron content and growth rate were conserved in *T. oceanica* (Table 1), suggesting that this species is better equipped to cope with iron limitation than *T. pseudonana*. Analysis of whole-cell protein relative abundances indicated that iron limitation leads to down-regulation of the photosynthetic electron transfer chain and light-harvesting components in *T. oceanica* (Fig. 4). This finding is consistent with the decreased chloroplast number observed when iron availability is limited (Fig. 3). Our analysis also revealed that the abundance of the chloroplast and mitochondrial ETCs components was conserved upon iron limitation in *T. pseudonana* (Fig. 4). These data suggest that, unlike *T. pseudonana*, *T. oceanica* favors respiratory processes over photosynthetic processes in conditions with moderate iron limitation. This hypothesis is in line with earlier findings in deeply-starved *T. oceanica* [3, 37] and other microalgae [44–46].

The differences observed between the two species in response to iron limitation cannot be attributed solely to variations in the abundance of molecular components of the photosynthetic chain. Rather, the cells adapt to control their energy use. As chloroplasts were distinctly positioned relative to mitochondria in the two species (Fig. 1, Fig. 2), with the relative positions of the organelles likely influencing their metabolic cross-talk (Fig. 5). We therefore propose that some of the dynamic differences reported here could be linked to the constitutive intracellular topology. Indeed, the adaptive structural differences between the two species are conserved upon iron limitation (Fig. 1, Fig.S2). Overall, it therefore appears that the phenotype is the outcome of both structural and metabolic differences.

In *T. pseudonana,* photosynthetic activity in the chloroplast is closely linked to mitochondrial activity (Fig. 5). This strong link is more than likely related to the specific subcellular organization of these cells, with the two organelles sharing a physical proximity that is not affected by iron limitation (Fig. 2, Fig. S2). The elongated shape of the mitochondria, a feature that is commonly observed in diatoms [47, 48] and other phototrophs [34], could be a determinant for this interaction. In cells grown with limited iron, the decrease in the total iron content tracks the decrease in total photosynthetic pigments content (Table 1). Both photosynthesis and respiration are severely decreased, and growth slows (Table 1). However, the energetic coupling between chloroplasts and mitochondria in iron-limited *T. pseudonana* cells (Fig. 5) could maintain a ‘high energy budget’ in a natural environment where iron is sufficiently abundant. The increase in expression of mitochondrial ATP synthase components (Fig. 4) could be part of this response, boosting energy exchanges by providing more ‘exchangeable’ mitochondrial-derived ATP [49]. In *T. pseudonana* cells grown in iron-poor medium, where the iron-demanding respiration activity is extensively diminished (Table 1), the exchanges between chloroplasts and mitochondria would only partially sustain the energy demands of photosynthesis. Therefore, the growth rate, which is mostly based on this energetic interaction, would ultimately collapse (Table 1).

In *T. oceanica,* unlike any other diatom species studied so far [47], mitochondria are spherical and have minimal physical contact with chloroplasts, with their shape creating geometric constraints on any potential interactions (Fig. 2, Fig. S2). As a result of the lack of physical contact, metabolic synergy is hampered, and energy exchanges between chloroplasts and mitochondria must occur by less efficient processes, such as diffusion or indirect exchanges (Fig. 5). In line with this hypothesis, our results indicated that photosynthesis depends less on mitochondrial function and thus on iron bioavailability in this species (Fig. 5, Table 1). In *T. oceanica*, limited iron availability triggers a reduction in the chloroplast compartment and its related components (Fig. 3, Fig. 4), whereas the rate of respiration remains almost constant, prioritizing maintenance of peak growth rates. The conservation of intracellular iron levels and iron-independent compensatory mechanisms, including constitutive expression of plastocyanin and reduced PSI to PSII ratios [26, 27] (Dataset S1), appear to provide further compensatory adaptations. Overall, the limited interdependence of chloroplasts and mitochondria, and their reduced reliance on iron for metabolic exchanges appear to reinforce these constitutive modifications. This functional acclimation strategy allows *T. oceanica* to maintain the relative homeostasis of its cellular energy processes, and, by reducing the energy demands of growth, is apparently sufficient to promote growth in the open-ocean environment [50, 51].

Based on our results, we conclude that: *i.* The photosynthesis rate is more dependent on mitochondrial function in *T. pseudonana* than in *T. oceanica*, and, *ii.* This interdependency is linked to iron availability in *T. pseudonana*, consistent with enhanced chloroplast-mitochondria synergy in this diatom. Because the iron concentrations used in our comparative analysis were chosen based on those observed in natural conditions, we believe that the conclusions of our in-laboratory study also apply in these natural oceanic environments.

## Materials and Methods

### Cultures, growth and establishing iron limitation

The coastal *Thalassiosira pseudonana* (Center for Culture of Marine Phytoplankton (CCMP) 1335) was originally isolated from Long Island Sound, NY, USA. *Thalassiosira oceanica* [CCMP 1005] was originally isolated from the Sargasso Sea. Cells were grown under controlled light (120 µmol.m-2.s-1; with a 12:12 photoperiod) at 18 °C, in artificial seawater (AQUIL [52]) supplemented with major nutrients, trace elements, and vitamins, in line with the recipe for L1 medium [53] published by the Culture Collection of Algae and Protozoa – CCAP (www.ccap.ac.uk). Silicate and copper concentrations were adapted based on the NCMA L1 media recipe (https://ncma.bigelow.org/). Culture media were supplemented with FeCl_3_ to produce iron-rich (113 nM) or iron-poor (18 nM) conditions. Cells were serially diluted until a constant growth rate was attained in each condition (∼7 generations, 3 passages), to ensure that a stable status was established [26, 32]. For all experiments, cells were harvested during the exponential growth phase, 2-3 h after initiating light illumination in the photoperiod.

### Iron quantification by inductively coupled plasma mass spectrometry (ICP-MS)

Cells were collected by centrifugation (15 min, 4000 x g, 20 °C). Pellets were washed three times in 0.3 M NaCl, 10 mM KCl, 20 mM HEPES pH 7.0 [30, 54]. Cell pellets were dehydrated to dryness before acid digestion at 90 °C for 4 h in 65% (w/v) nitric acid. Mineralized samples were analyzed on an iCAP RQ quadrupole mass spectrometer (Thermo Fisher Scientific) operating in helium collision mode. Magnesium (^25^Mg) and iron (mean of ^56^Fe and ^57^Fe) concentrations were determined by comparing signals from samples to standard curves produced using serial dilutions of a multi-element solution (ICP multi-element standard IV, Merck). Values were corrected thanks to an internal standard solution containing ^45^Sc and ^103^Rh, added online.

### Oxygen measurements

Oxygen was measured as described in [55]. Briefly, 700 μL of cell culture at 10 million cells/mL were maintained under gentle stirring at 18 °C in a WALZ KS-2500 water-jacketed chamber (Heinz Walz GmbH) paired with an FSO2-1 oxygen meter and optical microsensor (PyroScience GmbH). Before measurement, samples were bubbled with N_2_ for approximately 20 seconds to lower oxygen levels. NaHCO_3_ 5 mM was added to avoid carbon limitation during measurements. The gross oxygen evolution rate was calculated after ∼8 minutes of light exposure (120 μmol photons.m^-2^.s^-1^ PPFD). Oxygen consumption rate in the dark was measured just after extinction of the light source. This sequence of illumination was achieved thanks to a MINI-PAM-II controlled by WinControl-3 software (Heinz Walz GmbH).

### Pigment extraction and quantification

10-50 million cells were collected by centrifugation (4000 x g, 5 min, 4 °C). Protocols from previous studies [56–58] were adapted as follows: 1 mL of ethanol 96% was added to each tube, pellets were resuspended and incubated for 30 min at 4 °C to ensure complete solubilization. After incubation, samples were collected by centrifugation (1 min, 10000 x g). Total chlorophyll and fucoxanthin concentrations were calculated from absorbance measurements as follows:

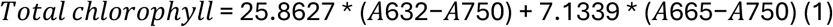

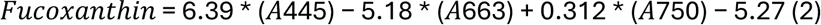

### Chlorophyll fluorescence measurements

Cell cultures were centrifuged (4000 x g, 5 min, 20 °C) and pellets were resuspended in culture media at the appropriate density (10 million cells/mL). The cell suspension (100 µL) was supplemented with NaHCO_3_ 5 mM to prevent carbon limitation during measurements. After dark acclimation (10 min), the maximum photochemical quantum yield of PSII (Fv/Fm) and the PSII operating efficiency (Φ _PSII_) were measured as described in [59]. An imaging fluorometer (Speedzen 3, JBeamBio, France) equipped with a highly sensitive and fast camera (Orca 4-Flash, Hamamatsu, Japan) was used. Tunable-intensity red light illumination was used to generate different actinic light irradiance levels (photon flux density – PPFD – from 0 to 250 μmol photons.m^−2^.s^−1^) and to produce a saturating pulse (3000 μmol photons.m^−2^.s^−1^). Chlorophyll fluorescence was induced by exposing the sample to blue light detection pulses. Experiments were performed at room temperature. Photosynthetic electron transport rate (ETR) through PSII was deduced from Φ _PSII_ using the following equation:

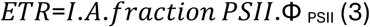

Where I is incident PPFD, A is the fraction of PPDF absorbed by the cells (assumed to be 0.84, i.e. 84%), fraction PSII is the fraction of absorbed light driving PSII photochemistry (assumed to be 0.5 for PSII, with the other half absorbed by PSI), and Φ _PSII_ is PSII operating efficiency [59].

Photosynthetic ETR was measured using the same setup, but adding respiratory inhibitors Antimycin A 1.2µM, [60] and salicylhydroxamic acid 250 µM [61]. These compounds respectively inhibit mitochondrial cytochrome c reductase (complex III) and alternative oxidase (AOX), thus targeting both the cyanide-sensitive and -insensitive respiratory pathways. [23].

### FIB-SEM sample preparation and data acquisition

Cells were harvested by centrifugation for 5 minutes at 2000 x g. Culture medium was eliminated, and pelleted cells were submitted to high-pressure freezing (HPM100, Leica) and freeze substitution (EM ASF2, Leica), as described previously [62, 63]. Focused ion beam (FIB) tomography was then performed on either a Zeiss NVision 40 or a Zeiss CrossBeam 550 microscope (Zeiss, Germany). The resin block containing the cells after substitution was fixed on a stub using carbon paste. It was then surface-abraded with a diamond knife in a microtome to obtain a clean, flat surface. Samples were metallized with 4 nm of platinum to avoid charging during observations. Inside the microscope, a second platinum layer (1–2 μm) was deposited locally on the area to be analyzed to mitigate possible curtaining artefacts. The sample was then scraped one slice at a time with a Ga+ ion beam (700-pA current at 30 kV), and each exposed layer was captured by SEM at 1.5 kV and a current of around 1 nA using an in-lens EsB backscatter detector. Individual SEM images – 1000-2000 frames, recorded to provide an isotropic voxel size of 8 nm – were combined to generate an image stack. Multiple cells from each culture were imaged.

### FIB-SEM data analysis

The FIB-SEM image stack was pre-processed to facilitate subsequent segmentation [34]. Fiji software was then used to crop individual cells from each pre-processed stack, thus reducing the number of frames to be analyzed. The segmentation method was adapted from previous studies [64] and was conducted using 3D Slicer 5.2.2 software tools (https://www.slicer.org/). Pixels were classified within each Region of Interest (ROI) as corresponding to cellular, subcellular, or organelle structures either manually or semi-automatically. For this step, a voxel volume of 1 (relative units) was set for image treatment and further corrected in line with the true voxel size (16 nm) during morphometric analysis. 3D representations of the segments were generated in STL format for subsequent smoothing and visual representation using MeshLab v2022.02–64-bit software (www.meshlab.net) and Paraview 5.11.0 software (https://www.paraview.org/). Morphometric analyses of volume and surface information for each ROI were performed using the Quantification/Segments statistics module in 3D Slicer Results and the Quality measures and computations module of MeshLab for comparison. The two methods returned comparable results and data were therefore averaged. Raw morphometric data can be found in supplementary file dataset_microscopy.xlsx.

### Proximity surface measurements

Simplified meshes of the 3D reconstructions (STL files) of chloroplasts and mitochondria were generated using MeshLab software’s Quadric Edge Collapse Decimation simplification algorithm. A reduction of 30% of the total number of mesh faces was applied to facilitate analysis by the minsurf.py Python script (from https://gitlab.com/clariaddy/mindist; [34]). This script was used to compute the total surface of a mesh present in a region within a threshold distance from a second mesh.

### Sample preparation for proteomics analysis

Cells (50 million) were harvested by centrifugation at 4000 x g, 4 °C, for 90 s to remove culture media. The pellet was then resuspended in 50 µL of Lyse reagent (PREOMICS iST sample preparation kit, www.preomics.com). Samples were incubated in line with the PREOMICS instructions and subjected to three cycles of sonication for 30 s to shear DNA. The resulting total protein extract was analyzed. Protein concentrations were determined using the PierceTM BCA protein assay kit (Thermo Scientific) according to the manufacturer’s instructions, using Bovine Serum Albumin to create a standard curve. For analyses, 20-35 μg of protein were digested and purified according to the instructions provided with the PREOMICS iST sample preparation kit.

### Mass spectrometry-based proteomics and data analysis

Approximately 500 ng of peptide mixture was analyzed by online nanoliquid chromatography (Ultimate 3000 RSLCnano) coupled to a Q-Exactive HF (Thermo Fisher Scientific). Peptides were concentrated on a precolumn (300 μm x 5 mm PepMap C18, Thermo Scientific) before separation on a 75 μm x 250 mm C18 column (Reprosil-Pur 120 C18-AQ, 1.9 μm; Dr. Maisch GmbH) by application of a 120-min gradient with the following profiles:; 0 – 10% solvent B (0.1% (v/v) formic acid in 80% (v/v) acetonitrile) over 8 min, 12 - 35% solvent B over 80 min, 36 - 44% solvent B over 10 min, 45-100% solvent B over 10 min, and finally 100% B for 10 min. Solvent A was 0.1% formic acid in water. Full-scan MS spectra were collected in a mass range of m/z 350 – 1300 Th in the Orbitrap at a resolution of 60’000 at m/z=200 Th after accumulation to an AGC (Automatic Gain Control) target value of 1e6 over a maximum injection time of 50 ms. In-source fragmentation was activated and set to 15 eV. The cycle time for the acquisition of MS/MS fragmentation scans was set to 2 s. Charge states accepted for MS/MS fragmentation were set to 3 - 8. Dynamic exclusion properties were set to n = 1 with an exclusion duration of 15 s. Stepped HCD (Higher energy Collision Dissociation) fragmentation (MS/MS) was performed with increasing normalized collision energy (27%). Mass spectra were acquired in the Orbitrap at a resolution of 30,000 at m/z=200 Th after accumulation to an AGC target value of 1e5 with an isolation window of m/z = 1.4 Th, and maximum injection time of 55 ms. Acquisition of MS and MS/MS data was controlled using Xcalibur (version 2.9; Thermo Fisher Scientific). Raw LC-MS/MS data were processed using MS-Fragger (v4.0) [65] integrated into FragPipe GUI (v21.1). Sequences were identified by searching against a database populated with the *Thalassiosira pseudonana* genome (taxonomy id 35128 downloaded from UniprotKB the 03/09/2024, 11’934 sequences), the *Thalassiosira oceanica* genome (taxonomy id 159749 downloaded from UniprotKB the 04/10/2024, 34’461 sequences) or the genome for the whole *Thalassiosira* genus (taxonomy id 35127 downloaded from UniprotKB the 14/10/2024, 47’418 sequences).

The latter database was used to retrieve a more complete list of proteins to compensate for the lack of a fully annotated *Thalassiosira pseudonana* database. An integrated contaminant database was used to filter out artefacts due to common contaminant proteins. For all searches, enzyme specificity was set to Trypsin/P (C-terminal cleavage of lysine and arginine) with up to two missed cleavages allowed. The minimum peptide length was set to 7 amino acids, and the maximum peptide mass was set to 6000 Da (∼55 amino acids). For peptide searches, fixed carbamidomethylation of cysteine and up to two variable modifications (methionine oxidation and protein N-terminal acetylation) were allowed per peptide. For database searches, precursor and fragment ion mass errors were set to 10 and 20 ppm, respectively. Match between runs was allowed within a retention time window of 0.5 min. A false discovery rate (FDR) of 1% for peptide spectrum matches (PSMs) and proteins was applied. Statistical analysis was performed using ProStaR software [66] based on the quantitative data obtained for the three biological replicates analyzed per condition. Proteins identified in the contaminant database, proteins identified by MS/MS in less than two replicates of one condition, and proteins quantified in less than two replicates of each condition were discarded. After log2 transformation, abundance values were normalized (variance stabilizing normalization) before imputing partially observed values (SLSA). Proteins for which no values were obtained in one condition were discarded. Differentially expressed individual proteins (volcano plots) were selected using a log2 (fold change) cut-off of 0.65 and a p-value cut-off of 0.006 (*T. pseudonana*) or 0.01 (*T. oceanica*). These parameters provided a FDR (False Discovery Rate) of less than 1% for both species according to the Benjamini-Hochberg estimator. When considering all the proteins linked to a single biological process (e.g. Calvin cycle, Krebs cycle), significant differences were selected based on a p-value cut-off of 0.01, regardless of the fold change (Dataset S1).

### Statistical treatments

All statistical analyses were performed using the Rstatix and Rsmic packages in RStudio. P-Values are indicated in the different figures and tables.

## Data availability

R Codes for data analyses (proteomic, physiology and proteomics) are available at https://zenodo.org/uploads/17748824#:~:text=10.5281/-,zenodo.17748824.

The mass spectrometry proteomics data have been deposited to the ProteomeXchange Consortium via the PRIDE [67] partner repository with the dataset identifier PXD071344.

## Acknowledgments

We acknowledge funding support from the European Research Council (ERC) through advanced grant ‘‘ChloroMito’’ (833184), and through the European Plankt-ON (101099192) project. We thank Benoit Gallet (Institute for Structural Biology, Grenoble France) for FIB-SEM sample preparation, Fabienne Devime (Plant and Cell Laboratory Grenoble, France) for technical support on the ICP-MS platform and the staff at EDyP proteomics platform (CEA Grenoble) for assistance with LC-MS/MS analyses. We thank Maighread Gallagher-Gambarelli for critical reading of the manuscript.

