## Supplemental Figures for "Chloroplast-mitochondria synergy modulates responses to iron limitation in two *Thalassiosira* diatom species"

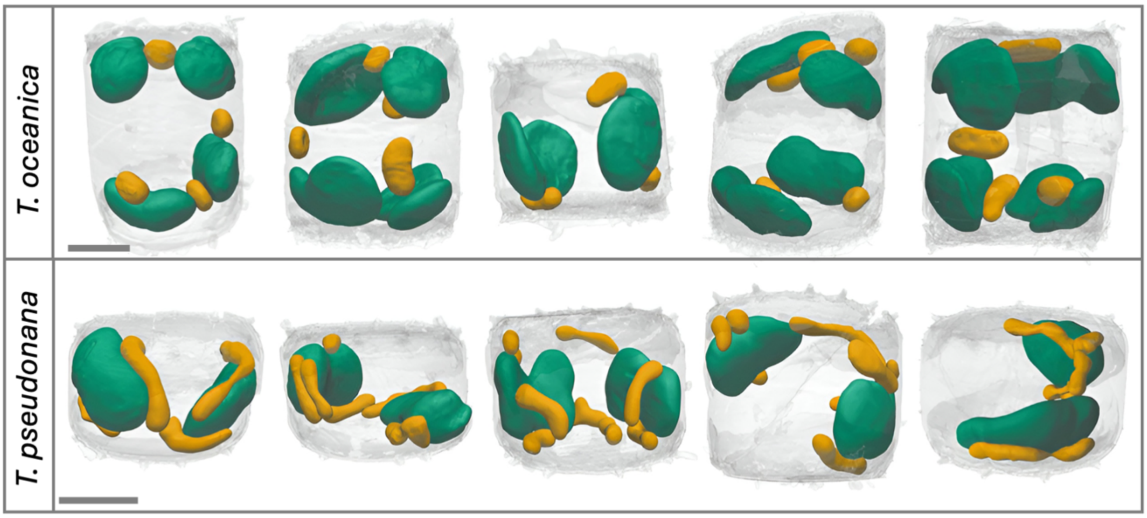


**Figure S1. 3D models of segmented cells of *T. oceanica* and *T. pseudonana*, showing chloroplast and mitochondria architecture under iron-rich conditions.** The cells were grown on iron-rich medium (see Material and methods). They were then imaged by FIB-SEM microscopy. Chloroplasts: green; mitochondria: yellow; cell frustule: gray. Scale bars = 2 µm. The 5 cells for each species presented here come from independent biological replicates.

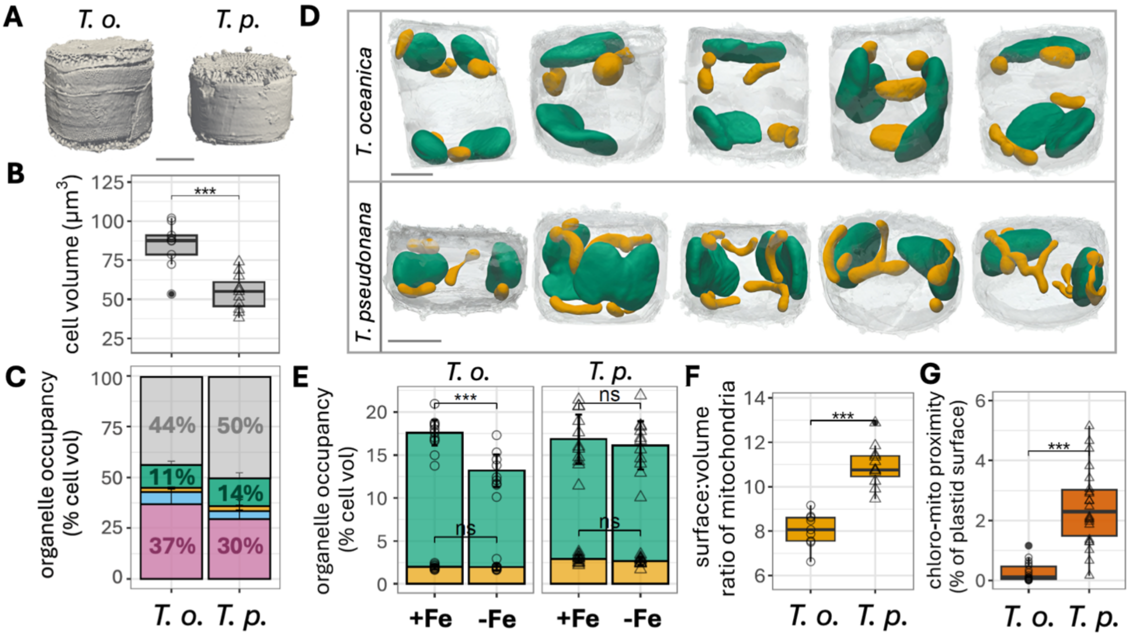


**Figure S2. Cell morphologies of *T. oceanica* and *T. pseudonana* under iron limitation.** T. oceanica and T. pseudonana cells were observed after growth in an iron-limited medium. The 3D models shown are representative of at least 9 cells from 2-3 independent biological replicates for each condition **(A)** 3D scan view of cell morphology. Voxel size = 16 nm. **(B)** Absolute cell volume, derived from quantitative analysis of the 3D models. **(C)** Volumes of the main subcellular compartments, derived from quantitative analysis of occupancy of the 3D models. Error bars: SD. Green: chloroplast; yellow: mitochondria; blue: nucleus; pink: vacuole; gray: other cell structures.**(D)** 3D models highlighting chloroplasts (green), mitochondria (yellow), and cell frustule (gray). **(E)** Total cell volume occupied by chloroplasts (green) and mitochondria (yellow), derived from quantitative analysis of the 3D models. Error bars: SD. For comparison, results from cells grown under Fe replete (+Fe) levels (cf. Fig. S1) are shown again. In *T. oceanica* specifically, the total chloroplast cell occupancy decreased from 15.6 ± 1.5 % to 11.2 ± 1.9 %. **(F)** Surface-to-volume ratio of total mitochondrial compartment in a single cell. **(G)** Total chloroplast surface at less than 30 nm from mitochondria (%). Mitochondria morphology and plastid-mitochondria proximity were unaffected upon iron limitation. **(A, D)** Scale bar: 2 µm. **(B, E, F, G)** T.o.: *T. oceanica*; open circles; n= 9 cells. T.p.: *T. pseudonana*; open triangles; n= 11 cells. Statistical analysis by paired Welch's t-test; ***: $p \leq 0.001$; ns: non-significant.

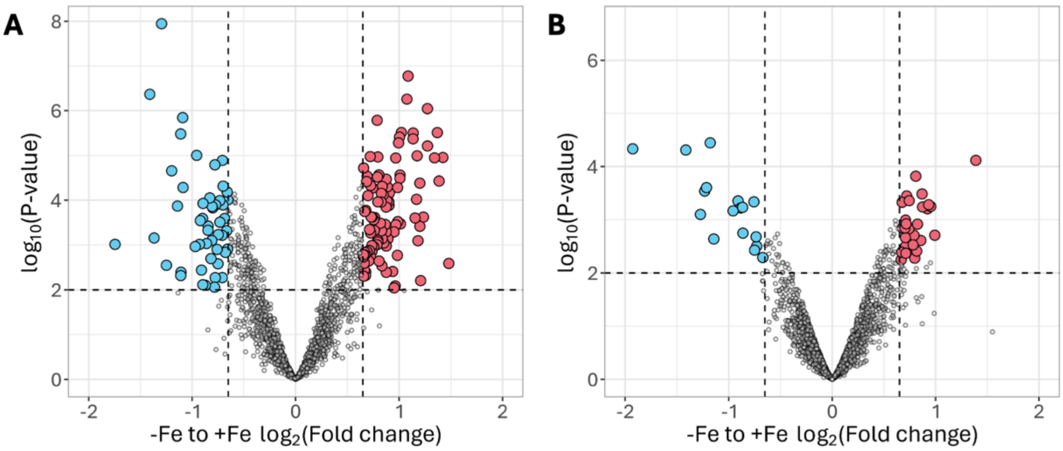


**Figure S3. Comparative Proteomic analysis of *Thalassiosira sp*. under iron limitation**. **(A)** *T. oceanica*. **(B)** *T. pseudonana*. Volcano plot displaying the differential abundance of proteins in the total proteomes analyzed by MS-based label-free quantitative proteomics. The volcano plot represents the $-\log_{10}$ (P-value), (Limma P-value, y axis) plotted against the $\log_2$FC (iron-limited vs. iron-rich, x axis) for each quantified protein. Blue and red dots represent significantly depleted and enriched proteins in iron-limited samples with respect to iron-rich samples, corresponding to a $\log_2$FC cut-off of 0.65 and a P-value cut-off of 0.006 for *T. pseudonana* and 0.01 for *T. oceanica* (thresholds as dashed lines, Benjamini–Hochberg False Discovery Rate < 1% respectively). Results are representative of 4 independent biological replicates. (See also Dataset S1).
